# Supervised non-negative matrix factorization on cell-free DNA fragmentomic features enhances early cancer detection

**DOI:** 10.1101/2024.12.20.629316

**Authors:** Trung Hieu Tran, Ngoc Tan Pham, Van Thien Chi Nguyen, Dac Ho Vo, Thi Hue Hanh Nguyen, Thi Trang Tran, Thanh Truong Tran, Truong Dang Huy Vo, Thi Huyen Dao, Huu Tam Phuc Nguyen, Thi Van Phan, Thi Minh Thi Ha, Thi Dieu Huong Ngo, Nhat Huy Tran, Nhat-Thang Tran, Thanh Quang Hoang, Viet Binh Nguyen, Van Cuong Le, Xuan Chung Nguyen, Thi Minh Phuong Nguyen, Van Hung Nguyen, Nu Thien Nhat Tran, Thi Ngoc Quynh Dang, Manh Hoang Tran, Phuc Nguyen Nguyen, Thi Anh Tuyet Pham, Duy Long Vo, Thuy Nguyen Doan, Viet Hai Nguyen, Quang Dat Tran, Quang Thong Dang, Le Minh Quoc Ho, Vu Tuan Anh Nguyen, Sao Trung Nguyen, Hoai-Nghia Nguyen, Le Son Tran, Hoa Giang, Minh-Duy Phan, Trong Hieu Nguyen

## Abstract

**Background:** Cell-free circulating DNA (cfDNA) fragments exhibit non-random patterns in their length (FLEN), end-motif (EM), and distance to nucleosome position (ND). While these cfDNA features have shown promise as inputs for machine learning and deep learning models in early cancer detection, most studies utilize them as raw inputs, overlooking the potential benefits of pre-processing to extract cancer-specific features. This study aims to enhance cancer detection accuracy by developing a novel approach to feature extraction from cfDNA fragmentomics.

**Methods:** We implemented a supervised non-negative matrix factorization (SNMF) algorithm to generate embedding vectors capturing cancer-specific signals within cfDNA fragmentomic features. These embeddings served as input for a machine learning model to classify cancer patients from healthy individuals.

**Results:** We validated our framework using two datasets: an in-house cohort of 431 cancer patients and 442 healthy individuals (dataset 1), and a published cohort comprising 90 hepatocellular carcinoma (HCC) patients and 103 individuals with cirrhosis or hepatitis B (dataset 2). In dataset 1, we achieved an AUC of 94% in pan-cancer detection. In dataset 2, our framework achieved an AUC of 100% for HCC vs healthy classification, 99% for HCC vs non-HCC patients classification, and 96% for identifying HCC patients among a mixed group of non-HCC patients and healthy donors.

**Conclusion:** This study demonstrates the efficiency of SNMF-transformed features in improving both pan-cancer detection and specific HCC detection. Our approach offers a significant advancement in leveraging cfDNA fragmentomics for early cancer detection, potentially enhancing diagnostic accuracy in clinical settings.

## Introduction

Cell-free circulating DNA (cfDNA) is a heterogeneous mixture of DNA fragments originating from various cellular sources upon apoptosis, necrosis, tumor-origin secretion in cancer patients or placenta-origin secretion in pregnant women. While a large proportion of cfDNA are derived from hematopoietic origin with minimal contribution from other sources, cfDNA fragments exhibit non-random patterns. Hence, observing an aberrant profile in an individual might suggest unusual epigenetic regulations of ongoing physiological conditions [1–2]. In the context of non-invasive prenatal testing (NIPT), fetal DNA constitutes approximately 10-15% of the total maternal cell-free DNA. Similarly, in early-stage cancer, tumor-derived cfDNA fragments account for a minor proportion, ranging from < 1% to 10% of the total plasma cfDNA in early and late stage cancer, respectively. Nevertheless, alterations in the cfDNA profile can still serve as effective predictive markers in numerous NIPT and early cancer screening studies [3–8].

In recent years, fragmentomics has emerged as prominent biomarkers of cfDNA. The term “fragmentomics” refers to the circulating DNA fragment size pattern, positioning and occupancy of nucleosomes, and fragment-end frequencies [3,9]. In healthy individuals, fragment length distributions have peaks corresponding to nucleosomes and chromatosomes at ∼143-147bp and ∼163-167bp [10], respectively. However, the peak of fragment length distribution in cancer patients commonly shifts toward the range of 90-150bp [11–13]. Profiling the landscape of fragment end motif in cfDNA is also an attractive approach. Observed cfDNA fragments in plasma samples undergo a fragmentation process that cleaves the genome at preferential genomic location [2,14]. Any alterations in genetic or epigenetic factors might cause aberrations in DNA endonuclease cleavage, leading to abnormal end-motif profiles [14]. Several 4bp end-motifs, namely CCCA, CCAG and CCTG, demonstrated significant differences in abundance when comparing cancer patients and non-cancer individuals [14,16]. Nucleosome footprints can also be discovered by fragmentation patterns in plasma cfDNA. A comprehensive map of all nucleosomes in the human genome has been constructed, and its utility in dissecting cell-type composition in healthy plasma samples has been demonstrated [1]. This proof-of-concept work was followed by a hypothesis suggesting that the presence of tumor-derived cfDNA fragments in a plasma sample could result in abnormal nucleosome occupancy and positioning. To investigate this hypothesis, the original window protection score (WPS) in [1] was refined (iwFAF) and combined with several nucleotide frequency features in [17]. Combined scores were then used as inputs to a random forest classifier, which achieved an AUC of 91% in pan-cancer detection.

Several studies have developed machine learning models for early cancer detection using fragmentomics features. In SPOT-MAS [18], fragmentomics features were combined with methylomics features in a multi-modal machine learning models. The model achieved a sensitivity of 72% at 97% specificity in early pan-cancer detection. Cristiano *et al.* [19] developed an XGBoost model using fragment length ratios in non-overlapping 5M-bp bins across the whole genome to classify cancer patients and control subjects in a cohort of 481 subjects. Fragment 4bp end-motif profile have been utilized for cancer detection in multiple forms, including raw frequencies, diversity measures (Shannon entropy) [15], low-rank 6-dimensional embeddings transformed by unsupervised non-negative matrix factorization [16], or concatenated sequences to be used in a language model [20]. Nucleosome-related features were also successfully employed in several machine learning models [17,21–22]. Notably, the distribution of distance from fragment to the nearest nucleosome was constructed, resulting in an M-shape distribution, which was subsequently used in a multinomial model to achieve an AUC of 89% in classifying ovarian cancer patients and healthy individuals [21]. Fragmentomic features derived from whole-genome bisulfite sequencing data (WGBS) have also demonstrated competitive performance in classifying early-stage cancers [22–24]. Nevertheless, most models have focused on individual aspects of fragmentomic features or used the features in their original form without any label-oriented feature engineering pre-processing steps.

Unsupervised non-negative matrix factorization (NMF) is a widely used technique for extracting informative features from high-dimensional data matrices [25–28]. A previous study has successfully applied NMF to transform fragment length distribution profiles into low-rank approximate embeddings, achieving an AUC of 96% when used as inputs for an SVM model [29]. Additionally, NMF deconvolution of end-motif distribution profiles has revealed multiple disease-related aberrant patterns [16]. To further exploit this feature transformation, we propose a supervised NMF (SNMF) algorithm by adding a classification-loss term to the original NMF loss function. This supervised approach is expected to further expand the advantages of guided feature learning by enabling the algorithm to incorporate the sample labels and extract informative data patterns that are distinguishable between the respective classes in the classifier loss function.

In this study, we implemented SNMF to transform fragment length distribution (FLEN), 4bp fragment end-motif (EM), and distribution of distance-to-nearest-nucleosome (ND) into 1D-embedding vectors. The SNMF embeddings derived from FLEN, EM, and ND were subsequently used as inputs for classification models. For simplicity, we refer to this entire process as the SNMF framework. To evaluate the performance of the SNMF framework, we conducted benchmarks using two datasets: an in-house cohort of 431 cancer patients and 442 healthy individuals (dataset 1), and a published cohort comprising 90 hepatocellular carcinoma (HCC) patients and 103 individuals with cirrhosis or hepatitis B (dataset 2, [13]). A schematic overview of the study is provided in Figure 1. We demonstrate the efficiency of SNMF-transformed features in improving both pan-cancer detection and specific HCC detection.

**Figure 1:**
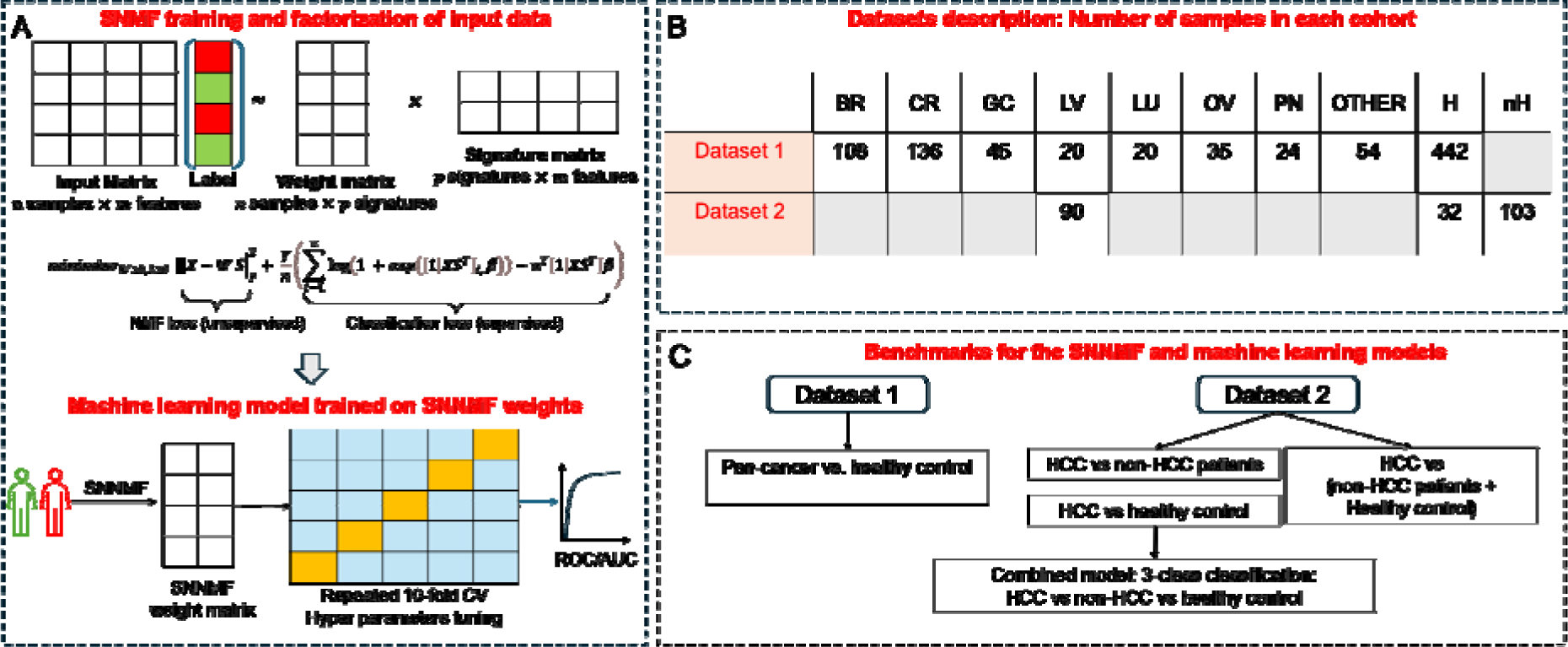
Schematic overview of the SNMF factorization and machine learning models. (A) A general framework of SNMF. The SNMF algorithm combining a NMF loss function and a classification loss function was used to generate low-dimensional embeddings (SNMF weight matrix), which were used to train machine learning classification models in a repeated 10-fold cross validation. (B) Number of patients in each cancer type, healthy individuals (H) and non-cancer patients (nH). Cancer types includ BR - breast, CR - colorectal, GC - gastric, LV - liver, LU - lung, OV - ovarian and PN - pancreatic. Other cancer types, including laryngeal, kidney, head and neck, endometrial, esophageal, bileduct and cervical cancers, were group to a class “other” due to their small sample sizes. (C) Overview of benchmarks conducted in this study.

## Results

### SNMF reveals cancer-specific FLEN, EM and ND signatures

To demonstrate the potential of extracting cancer-specific signals by SNMF in dataset 1, we first transformed FLEN and ND into 2-dimensional embeddings, while EM was transformed into a 6-dimensional embedding. Such low-dimension embeddings were chosen to facilitate visualization. We used all healthy donors and cancer patient samples in the training set of dataset 1 to train the SNMF.

Signature matrices obtained from this training were used to transform the held-out test set. Weight associated with each signature were normalized to have a total summation of 1 within each sample. The input features and their factorizations are illustrated in Figure 2. The fragment length distribution associated with *signature 2* exhibits a left-ward shift with multiple smaller peaks within the range of 100-150bp (Figure 2B), while *signature 1* exhibits a more right-ward shift with peaks at 160-167bp. Smaller peaks observed at short fragment lengths (<150bp) have been reported, and may reflect the abundance of tumor-derived cfDNA [11–13,18]. Additionally, *weight 2* corresponding to *signature 2* is significantly higher in cancer compared to healthy samples (t-test p-value < 0.05), suggesting that signature 2 represents the tumor-derived source of cfDNA in the samples (Figure 2C). A similar observation was obtained for ND features, where the *weight 2* corresponding to *signature 2* was also significantly associated with the cancer label (Figure 2E, F). For EM features, we used SNMF to construct a 6-dimension embedding of all 4bp end motifs as previously done by NMF [16] (Figure 2J). Consistently, among 6 signatures, we observed that normalized weights associated with *signatures 3* and *4* showed most significant differences between cancer and healthy samples (Figure 2H, I). These results demonstrate the ability of SNMF to extract cancer-specific signals from all three types of input features. In practice, we chose a higher value for the number of signatures *p* to obtain a better approximation of the input feature matrix (Supplementary Figure 3, Materials and methods).

**Figure 2:**
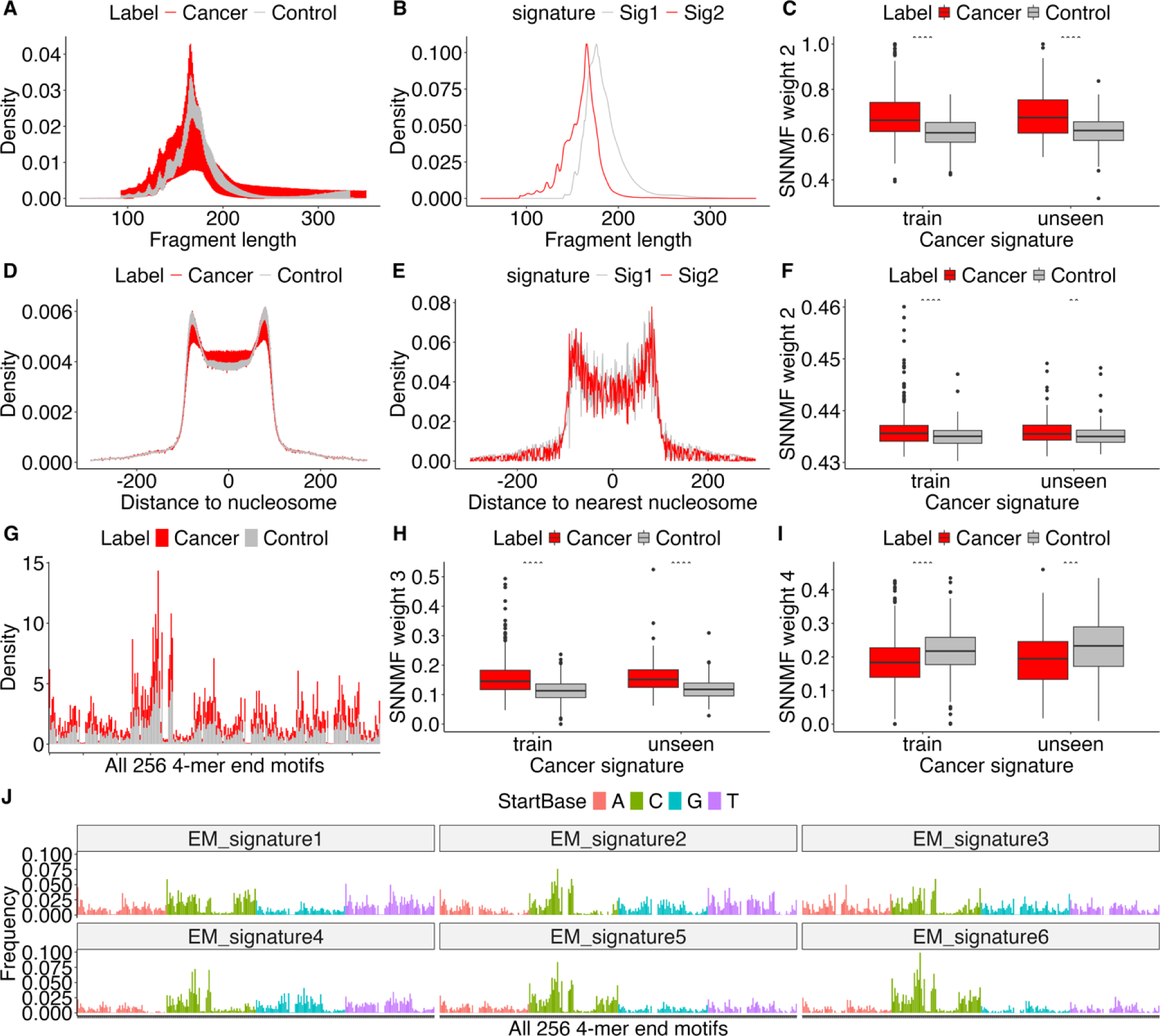
Cancer-specific signatures from FLEN (A-C), ND (D-F) and EM (G-J) are revealed by SNMF transformations. FLEN and ND were transformed to a 2-dimensional space while EM was transformed to a 6-dimensional space. (A) FLEN in cancer and healthy samples. (B) Two signatures extracted from FLEN by SNMF. The lower-peak signal (red) was assumed to be a cancer-related signal. (C) Weight corresponding to the FLEN lower-peak signal (red) is significantly associated with cancer samples. (D) ND in cancer and healthy samples. (E) Two signatures extracted from ND by SNMF, where signature 2 was assumed to be a cancer-related signal. (F) Weight corresponding to the signature 2 in (E) is significantly different between cancer and healthy samples. (G) Raw frequencies of all 256 4bp end motifs in cancer and healthy samples. (J) The 6 signatures of 256 4bp end motif extracted from (G). (H) and (I) Weights corresponding to signatures 3 and 4 in (J) are significantly different between cancer and healthy samples. All statistics tests were done with t-test.

### Detecting pan-cancer using SNMF framework

To evaluate cancer detection performance, we applied the SNMF framework to dataset 1. We trained the SNMF on a training set of 608 samples (298 healthy individuals, 310 cancer patients) and validated on a held-out testing set of 265 samples (133 healthy individuals, 132 cancer patients). Initially, the SNMF framework was trained independently for each feature, and the resulting models achieved the AUCs of 74%, 85% and 87% for ND, FLEN and EM features, respectively (Figure 3A). To assert whether combining the embeddings improves the performance, we concatenated embeddings from ND, FLEN and EM, and re-train machine learning classification models. Indeed, we achieved an enhanced performance with an AUC of 94%, and a sensitivity of 80% at 93% specificity in classifying cancer patients from healthy control samples (Figure 3) on the held-out test set of dataset 1. Stratifying by individual cancer types, the model demonstrated the ability to detect a diverse range of cancers effectively (Figure 3B). The common cancer types (colorectal, breast, gastric, liver and lung) exhibited a higher sensitivity of 90% compared to the rare cancer types (ovarian, pancreatic, cervical, esophageal, endometrial, head and neck, kidney, biliary tract and laryngeal), of which the model achieved a sensitivity of 63%. The lower sensitivity of rare cancer types might reflect the diversity of cancer types in this category and the scarcity of samples within each type in dataset 1.

**Figure 3:**
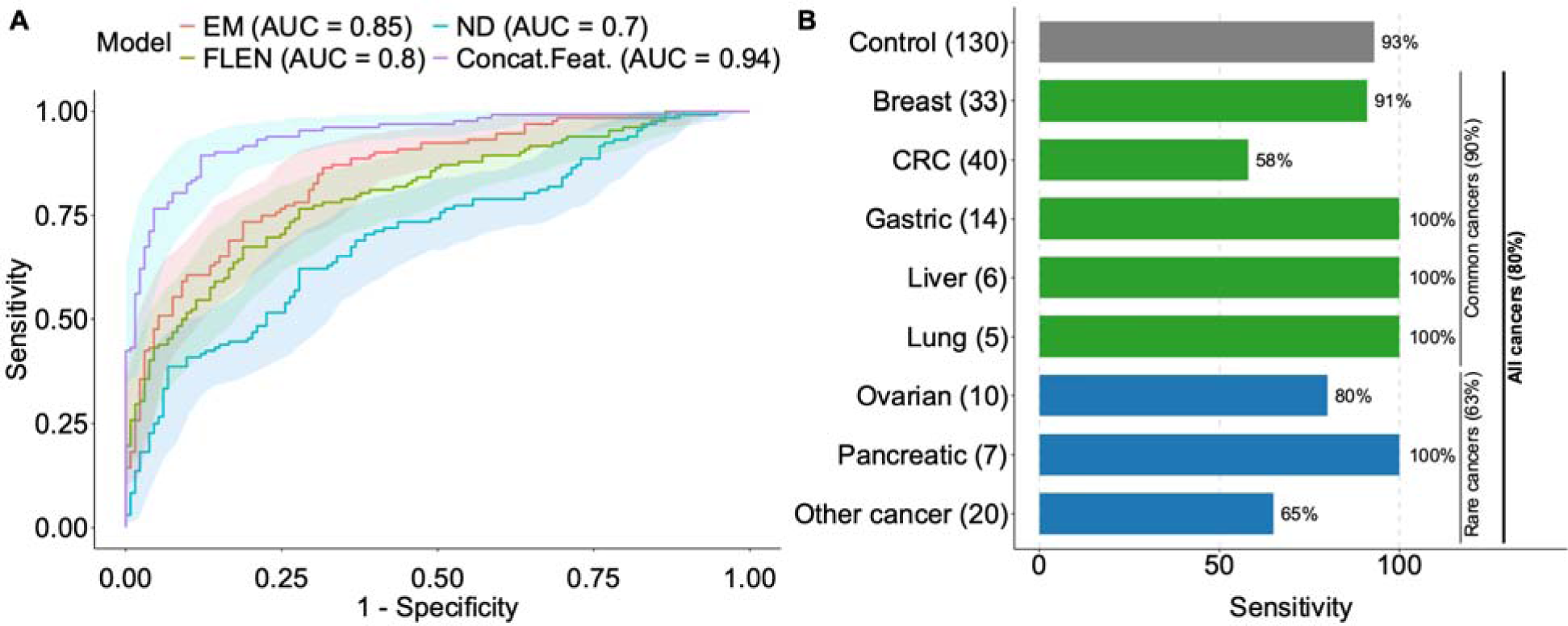
Pan-cancer detection performance of the SNMF framework. (A) ROC curve and AUC of models trained on SNMF-transformed EM, ND, FLEN and an embedding-concatenated model. (B) Specificit (class Control, 93%) and sensitivities of the embedding-concatenated model from the SNMF framework on each cancer type on the held-out test set of dataset 1. The combined sensitivities of all cancers, common cancers and rare cancers are also provided in brackets. The number of samples in each class is shown in brackets next to the class label. Other cancer includes laryngeal (n=1), biliary tract (n=2), kidney (n=4), head and neck (n=6), endometrial (n=7), esophageal (n=14), cervical (n=20).

### SNMF effectively distinguishes HCC from non-HCC patients and healthy individuals

We next evaluated the ability of the SNMF framework to classify HCC patients, patients with non-cancerous liver diseases (hepatitis B and cirrhosis), and healthy individuals using dataset 2. This evaluation is crucial for demonstrating that the method could maintain a low false-positive rate in a high-risk population, where signals from non-cancerous liver diseases can be misinterpreted as cancer signals.

Such differentiation is of particular importance for the reliability and clinical utility of an early cancer detection test [32–33]. We first trained the SNMF framework to perform two binary classification tasks: HCC patients vs healthy individuals (Figure 4A) and HCC patients vs non-HCC patients (Figure 4B). For the HCC vs. healthy classification, our concatenated model achieved an average AUC of 100% (Figure 4A). In the HCC vs non-HCC classification, we achieved the performance at an AUC of 99% with the EM model and 98% with the concatenated model (Figure 4B). Next, we train the framework to classify HCC patients from a combined group of non-HCC patients and healthy donors. In this binary classification setup, non-HCC patients and healthy donors were treated as a single class. The highest performance was achieved with the concatenated model with an AUC of 96% (Figure 4C). To classify all three classes (HCC, non-HCC, healthy), we developed a stacked model by combining the HCC vs non-HCC model and the HCC vs healthy model to derive a 3-class classification model (Figure 4D). This stacked model achieved an accuracy of 99%, 91% and 94% for the HCC, healthy, and non-HCC class, respectively (Figure 4D). These validations demonstrate the robust ability of the SNMF framework to differentiate HCC patients from non-HCC patients and healthy individuals. The high AUC values across various classification tasks and the successful implementation of a stacked model for multi-class classification underscore the potential of this approach for early HCC detection and differentiation from other liver conditions in clinical settings.

**Figure 4:**
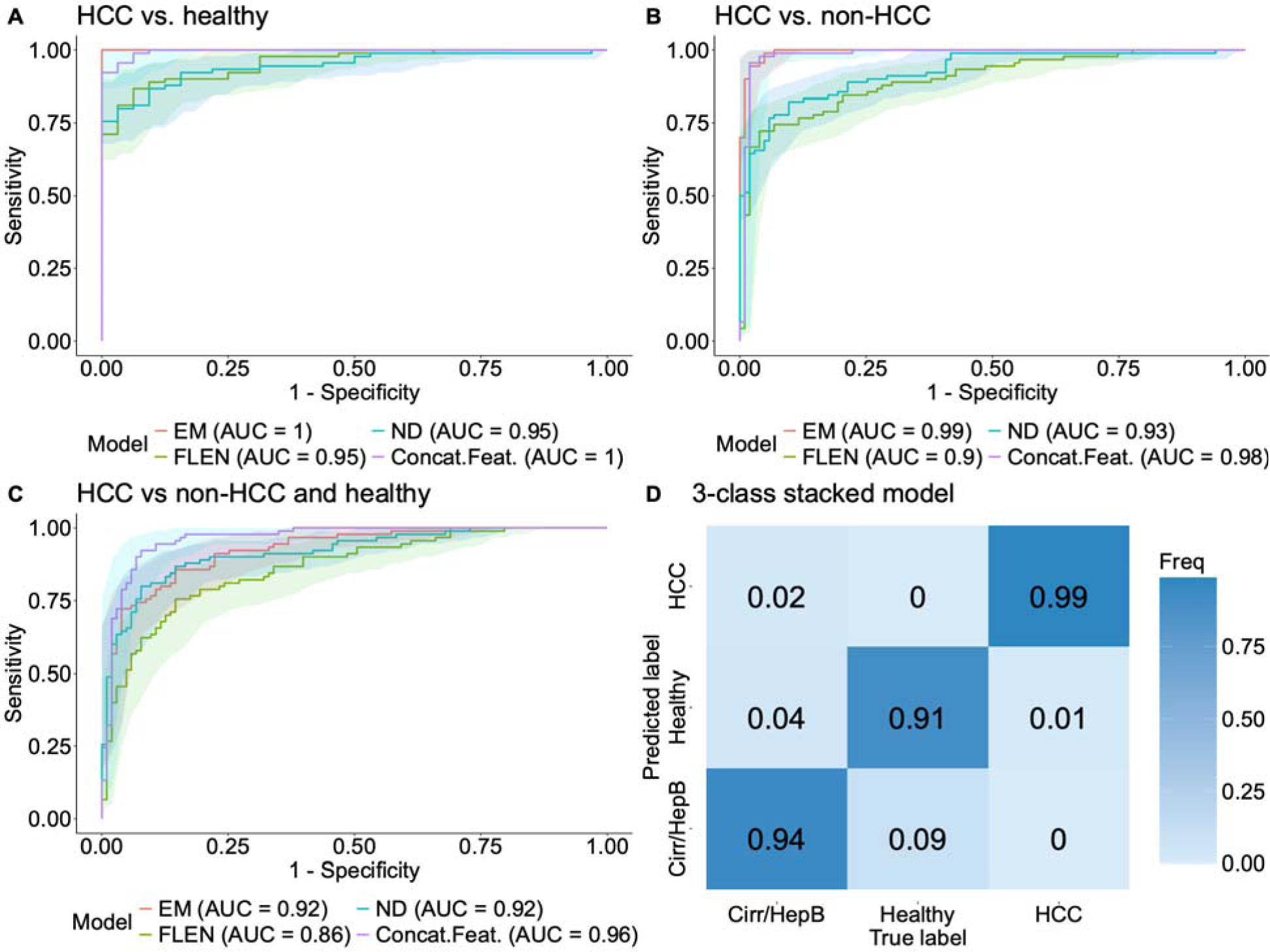
The SNMF framework demonstrated high classification performance on dataset 2. (A) ROC curves and AUCs of EM, FLEN, ND and concatenated models for classifying HCC patients vs health individuals. (B) ROC curves and AUCs of EM, FLEN, ND and concatenated models for classifying HCC patients vs non-HCC patients. (C) ROC curves and AUCs of EM, FLEN, ND and concatenated models for classifying HCC patients vs a combined class of healthy individuals and non-HCC patients. (D) Confusion matrix obtained from the stacked model for 3-class classification.

## Materials and methods

### Datasets used in this study

This study used two distinct datasets. Dataset 1 (in-house cohort) comprised of 431 healthy individuals and 442 cancer patients. The cancer group included laryngeal (n=1), biliary tract (n=2), kidney (n=4), head and neck (n=6), endometrial (n=7), esophageal (n=14), cervical (n=20), liver (n=20), lung (n=20), pancreatic (n=24), ovarian (n=35), gastric (n=45), breast (n=108), and colorectal (n=136). All samples underwent whole-genome sequencing at a low depth of coverage (∼0.5x). We randomly partitioned this dataset, allocating 70% for training the SNMF and subsequent machine learning models, and reserving 30% as a held-out test set for final model evaluation. Detailed metadata for this cohort are provided in Supplementary Table S1. Dataset 2 [13] was obtained from FinaleDB [32] and comprised of 32 healthy control samples, 36 patients with cirrhosis, 67 patients with hepatitis B and 90 liver cancer patients (HCC).

### Patient enrollment for dataset 1

This study recruited 431 healthy individuals and 442 cancer patients. All cancer patients were confirmed to have one of the cancers analyzed in this study. Cancer patients were confirmed to have cancer by abnormal imaging examination and subsequent tissue biopsy confirmation of malignancy. All healthy subjects were confirmed to have no history of cancer at the time of enrollment. They were followed up at 6 months and 1 year after enrollment to ensure that they did not develop cancer. Study subjects were recruited from the University of Medicine and Pharmacy, Thu Duc City Hospital, University of Medicine and Pharmacy – Hue University, University Medical Center HCM, Thai Nguyen Central Hospital, Nghe An Oncology Hospital, Ha Tinh General Hospital, Nghe An Maternity-Pediatric Hospital, 108 Military Central Hospital, Military Hospital 175, Thong Nhat Hospital, Binh Dan Hospital, Tuyen Quang Provincial General Hospital, Medical Genetics Institute in Ho Chi Minh City, Vietnam, National Cancer Hospital and Hanoi Medical University in Hanoi from May 2019 to June 2024.

Written informed consent was obtained from each participant in accordance with the Declaration of Helsinki. This study was approved by the Ethics Committee of the Medic Medical Center, University of Medicine and Pharmacy (K-Discovery A, approval number 192/HDDD-DHYD) and Medical Genetics Institute (K-Discovery B, approval number 04/2024CT-VDTYH), Ho Chi Minh City, Vietnam. All cancer patients were treatment-naïve at the time of blood sample collection.

### Isolation of cfDNA

Each participant provided 10 ml of peripheral blood, which was collected in a Cell-Free DNA BCT tube (Streck, USA). The samples underwent two rounds of centrifugation—first at 2,000 × g for 10 minutes, followed by 16,000 × g for 10 minutes—to separate the plasma from blood cells. The plasma fraction was collected and aliquoted into 1 ml portions. The samples were then stored at −80°C. cfDNA extraction was performed on the plasma aliquots using the MagMAX Cell-Free DNA Isolation Kit (ThermoFisher, USA) according to the manufacturer’s protocol. The resulting cfDNA was quantified using the QuantiFluor dsDNA system (Promega, USA).

### Library preparation and sequencing

The library preparation was done using NEBNext Ultra II DNA Library Prep Kit from New England BioLabs (Ipswich, MA, USA), adding adapters to the cfDNA samples. DNA concentrations were measured using the QuantiFlour dsDNA system (Promega, USA). Samples were sequenced on the DNBSEQ-G400 DNA sequencing system (MGI Tech, Shenzhen, China) to generate sequencing data with 100-bp paired-end reads, at a sequencing depth of 15 million reads.

### Bioinformatics pipelines

For dataset 1, all FASTQ files were examined by FastQC v. 0.11.9 and MultiQC v. 1.12. Sequences were trimmed by Trimmomatic (v. 0.39, [33]) with the input parameters (CROP:50 ILLUMINACLIP:TruSeq3-PE-2:2:30:10 LEADING:3 TRAILING:3 SLIDINGWINDOW:4:15 MINLEN:36). Trimmed sequences were then aligned by bwa-mem (v. 0.7.17, [34]) to the human genome version hg38. Duplicated sequences were marked by picard MarkDuplicates with default parameters (v 3.2.0, https://gatk.broadinstitute.org/hc/en-us/articles/360037052812-MarkDuplicates-Picard). Samtools (v 1.13, [35]) was utilized to work with SAM/BAM alignment files. For dataset 2, all data were pre-processed in FinaleDB and we obtained tab-separated files containing chromosome, starting and ending position for all DNA fragments. To generate FLEN, EM, and ND features, we used our in-house scripts written in python as described in the github repository (Code availability).

### Construction of the SNMF and its downstream machine learning models

#### General framework

We constructed a supervised non-negative matrix factorization algorithm to transform the fragment length distribution (FLEN), 4bp end-motif distribution (EM) and distribution of distance to nearest nucleosome (ND) features to low-dimensional approximations. By adding a logistic regression binary classification loss into the original non-negative matrix factorization (NMF) loss, we aim to guide the factorization process to better preserve cancer-specific signatures within the extracted features. Each input feature matrix (*n samples* x *m features*) was factorized into a *weight* matrix (*n* x *p*) and a *signature* matrix (*p* x *m*). After obtaining the embeddings, they were used as inputs for a series of machine learning models including logistic regression, suppport vector machines, random forest and XGboost. Details are given in the following two sections.

### Supervised non-negative matrix factorization algorithms

To implement the SNMF, we followed the implementation outlined in [36]. For simplicity, we denote by *X* the input feature matrix (FLEN, EM or ND) of shape *n* samples x *m* features. The main goal is to decompose the input matrix *X* into *W* (<u>w</u>eight) and *S* (signature), 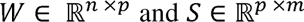, where *p* is a pre-defined number of transformed dimensions and *p* should not be large, as *SNMF* can also be considered as a low-dimensional reduction technique. The matrix *S* represents a set of *p* most dominating sources of signatures existing in the input data *X* and *W* is a low-dimensional approximation of the input matrix *X*. To find *W* and *X* unsupervisedly, we solve the following optimization problem

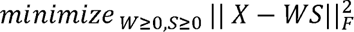

where ||.|| denotes the Frobenius matrix norm. The minimization problem subjects to the condition that *W* and *S* are both positive.

To turn the above unsupervised minization problem into a supervised problem, we add a classification loss function. In this work, we chose the loss function of a logistic regression (logit model),

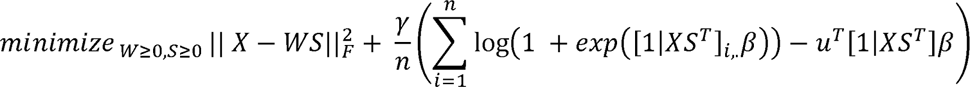

where y > 0 is a regularization parameter to balance betweeen the unsupervised NMF loss and the classificatio loss. The vector-value *u* encodes the information on samples’ labels, which make the optimization problem supervised.

To solve this supervised problem, we apply the same Majorization – Minimization algorithm (MM) ([36], [37]). In short, the MM algorithm defines a surrogate function which majorizes the objective function. To majorize a given function *f(x)*, we define a function *g_a_(x)*. Such surrogate functions should satisfy: (1) 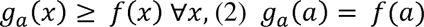 and (3) the function *g_a_* should be *easy* to minimize than the original function *f*. This construction yields an iterative update rule 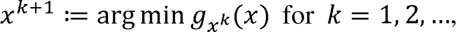

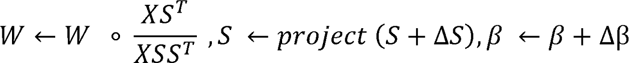

where *project* denotes the gradient based update projection on *R_+_*. The sequence {*x^k^*}*_k=12,…_* is proved to converged to a solution. In practice, the sequence converges to a minimizer of *f*. However, such a minimizer is not guaranteed to be a global minimizer but rather be a local minimum.

Since solving SNMF with MM is an iterative algorithm, initialization plays a crucial role. We benchmarked multiple approaches to initializations, such as Nonnegative Double Singular Value Decomposition (NNDSVD), NNDSVD with zeros filled with the average of X, NNDSVD with zeros filled with small random, and random initialization ([38]). We also performed grid-search for *p*, number of low dimensions, from 100 to 300. Tuning this parameter poses a trade-off between minimizing the approximation error and managing the complexity of the resulting high-dimensional feature space (Supplementary Figure 3). By selecting a higher *p*, we balanced the need for a more accurate approximation of the input feature with the curse of high dimensionality. Since SNMF is an iterative solver, we evaluated various initialization methods and values for the low-dimensional approximation parameter *p* to find the best combination that minimizes reconstruction loss. In practice, for each feature, a combination of *p* and initialization method, which achieves the lowest reconstruction loss, is chosen.

### Machine learning classification models on SNMF embeddings

To perform classification on the SNMF embeddings, we implemented a range of machine learning models, including logistic regression, support vector machines, random forests, and XGBoost. Each model was trained and evaluated separately for each feature type. Additionally, we investigated whether concatenating embeddings derived from three distinct feature types (FLEN, EM, and ND) could enhance classification performance. Model training and evaluation were conducted using repeated 10-fold cross-validation. We focused on achieving a minimum specificity of 95%, with optimal sensitivity under this constraint. We also benchmarked individual feature models (FLEN, EM, ND) and a concatenated feature model, in which embeddings are concatenating into a 1D vector. Samples were assigned to training and testing sets in a way that the number of cancer patients in each set are balanced. All models were implemented using custom Python scripts and the Scikit-learn library [38].

## Discussion

The application of fragmentomic features (FLEN, EM, and ND) has become one of the most popular approaches in cancer detection, demonstrating remarkable success across various applications [10,16–18,22–24]. However, most existing methodologies use the original forms of these features, often neglecting the potential benefits of feature engineering or transformation. Such techniques might hold significant promise for enhancing classification performance. It is noteworthy that the FLEN, EM and ND features are comprised of signals from differential sources, contributing to the overall mixture of patterns. For instance, tumor-derived fragments might have shorter fragment lengths [2], [11], [19], aberrant motifs due to unusual cleavage of DNA fragmentation [15–16] and variations in nucleosome depleted regions [1], [17], [21]. These observations highlight the need for a feature engineering method that is capable of decomposing signals into various sources. One such method, the unsupervised non-negative matrix factorization (NMF) has been used to transform FLEN [29] and EM features [16] and extract cancer-related signals. In this study, we proposed an application of the supervised non-negative matrix factorization algorithm to transform fragmentomic features in a supervised manner. The transformed vectors were then used as inputs to a range of machine learning models for a binary classification task (ie. cancer vs healthy). Although SNMF has been widely applied across diverse fields [36], its application to fragmentomic features for a cancer detection task has not been explored. We believe that this method, which is guided toward the differences between specific cancer vs healthy signals, could offer a greater utility and enhance classification performance.

We first validated our SNMF framework on our in-house dataset (dataset 1). Our best model, which combined embeddings from FLEN, EM, and ND features, achieved an AUC of 94% in a held-out test set. The model demonstrated stable performance in predicting common and rare cancer types. For common cancer types (colorectal, breast, gastric, liver, and lung), the model achieved a higher sensitivity of 90%, while for rare cancer types, it showed a sensitivity of 63%, all while maintaining a specificity of 93%. This performance was achieved through a stepwise approach. Initially, the SNMF framework was trained independently for each feature type, resulting in AUCs of 74%, 85%, and 87% for ND, FLEN, and EM features, respectively. The enhanced performance (AUC of 94%, sensitivity 80% at specificity 93%) was achieved by concatenating embeddings from all three feature types, demonstrating the synergistic effect of combining these fragmentomic features. With this model, we achieved comparable performance to other studies, such as DELFI [19] (specificity 98%, sensitivity 78%) and SPOT-MAS [18] (specificity 97%, sensitivity 72% for common cancers). Our findings demonstrated that the SNMF transformation and its downstream machine learning classification algorithms are robust and effective in classifying cancer patients and healthy individuals across a diverse range of cancer types. This is particularly noteworthy for clinical utility given the heterogeneity of the cancer patients.

Our second validation investigated whether the framework could distinguish HCC patients from patients with other liver diseases (non-HCC), including cirrhosis and hepatitis B, and healthy donors. We built and evaluated 3 approaches: HCC vs. non-HCC patients, HCC vs. healthy donors, and HCC vs. non-HCC and healthy donors. Overall, the SNMF framework demonstrated excellent performance in all tasks, achieving AUC of 100%, 98% and 96%, respectively.

Our study has several limitations. Firstly, while dataset 1 provided a sufficient sample size for pan-cancer model training and evaluation, the uneven distribution of cancer types, dominated by breast and colorectal cancers, may bias the model towards these prevalent types. This imbalance could potentially reduce the model performance in detecting less represented cancer types, highlighting the need for careful curation of training datasets. Secondly, the confounding effects introduced by the variations in tumor-derived cfDNA in each sample across cancer types and stages were not accounted for in this study. Thirdly, we only tried to incorporate the loss function from a logit model into our SNMF framework. Exploring other classification loss functions, such as Hinge loss or linear discriminant analysis loss, could potentially enhance both feature extraction and classification. However, other loss functions might present challenges in solving the combined SNMF loss function with a practical optimization algorithm. Finally, our approach did not provide an estimate of tumor fraction. Future work should focus on developing a method to determine the tumor fraction in a cfDNA sample based on the relative contribution of each source of signals after SNMF decomposition.

In conclusion, we have implemented a supervised non-negative matrix factorization algorithm to decompose FLEN, EM and ND to obtain low-dimensional representations, which were guided to highlight the difference between the two classes in a given training set. Embeddings were then used as inputs to machine classification models to obtain final predictions. Our approach is not only capable of detecting pan-cancer, including a wide range of common and rare cancer types but also effectively distinguishes HCC from other non-cancerous liver diseases commonly found in high-risk populations. The SNMF framework offers a significant advancement in leveraging cfDNA fragmentomics for early cancer detection, potentially enhancing diagnostic accuracy in clinical settings.

## Code availability

All code used for the building of SNMF and its machine learning models are available at https://github.com/hieunguyen4193/gs_snmf.

## Declarations

### Ethics approval and consent to participate

This study was approved by the Ethics Committee of the Medic Medical Center, University of Medicine and Pharmacy and Medical Genetics Institute, Ho Chi Minh city, Vietnam. Written informed consent was obtained from each participant in accordance with the Declaration of Helsinki.

### Consent for publication

Not applicable.

### Availability of data and materials

Analytic data are available on request to the corresponding authors (THN, MDP). Raw FASTQ data are not available due to ethical and regulatory restrictions. We confirm that this does not alter our adherence to Cancer Investigation policies on sharing data and materials. Dataset 2 is available in the FinaleDB database [32].

### Authors’ contributions

Conceptualization: THN, MDP, HG, LST

Data analysis and machine learning model implementation: THT, NTP, THN

Wetlab experiments: VTCN, DHV, THHN, TTT, TTT, TDHV, THD, HTPN

Patient consultancy and screening: TMTH, XLT, NHT, TQH, VBN, VCL, XCN, TMPN, VHN, NTNT, TNQD, MHT, PNN, NTT, TATP, DLV, TND, VHN, QDT, QTD, VTAN, LMQH

Writing-original draft: THN, THT

Writing-review and editing: THN, MDP, HG, LST

---

**Supplementary Figure 1.**
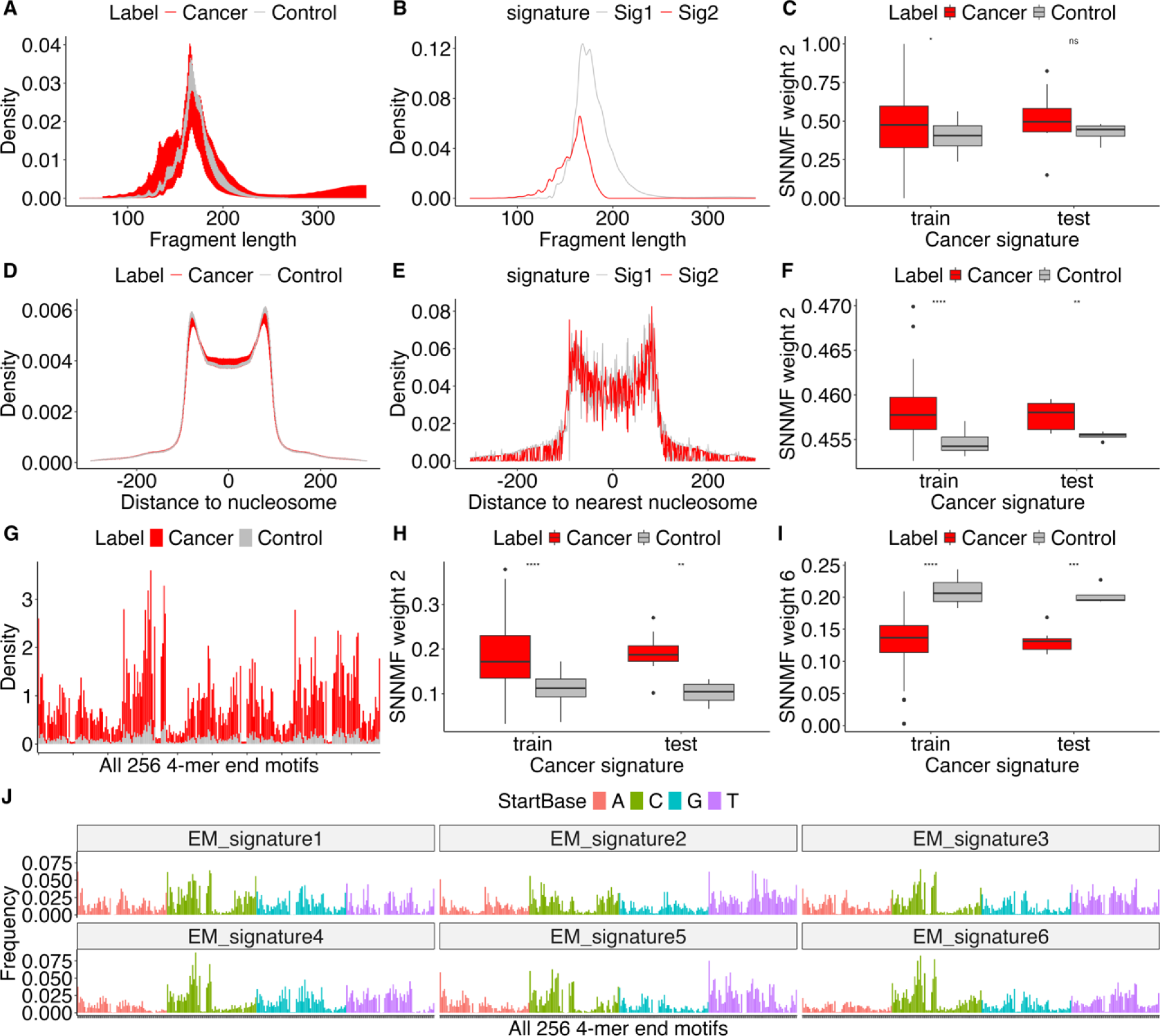
Cancer-specific signatures from FLEN (A-C), ND (D-F) and EM (G-J) are revealed by SNMF transformations when trasnforming held-out testing set of dataset 2. FLEN and ND were transformed to a 2-dimensional space while EM was transformed to a 6-dimensional space. (A) Fragment length distributions in cancer and healthy samples. (B) Two signatures extracted from (A) by SNMF. The lower-peak signal (red) was assumed to be cancer-related signal (C) Weights associated to the lower-peak signal (red) is higher in cancer samples. (D) Raw distributions of distances to nearest nucleosome (ND) in cancer and healthy samples. (E) Two signatures extracted from (D) by SNMF. (F) Weights associated to the two signals in (E) are different between cancer and healthy samples. (G) Raw frequencies of all 256 4bp end motif in cancer and healthy samples. (J) The 6-profile of 256 4bp end motif extracted from (G). (H) and (I) Weights associated to the 2^nd^ and 6^th^ signatures in (J) are different between cancer and healthy samples.

**Supplementary Figure 2.**
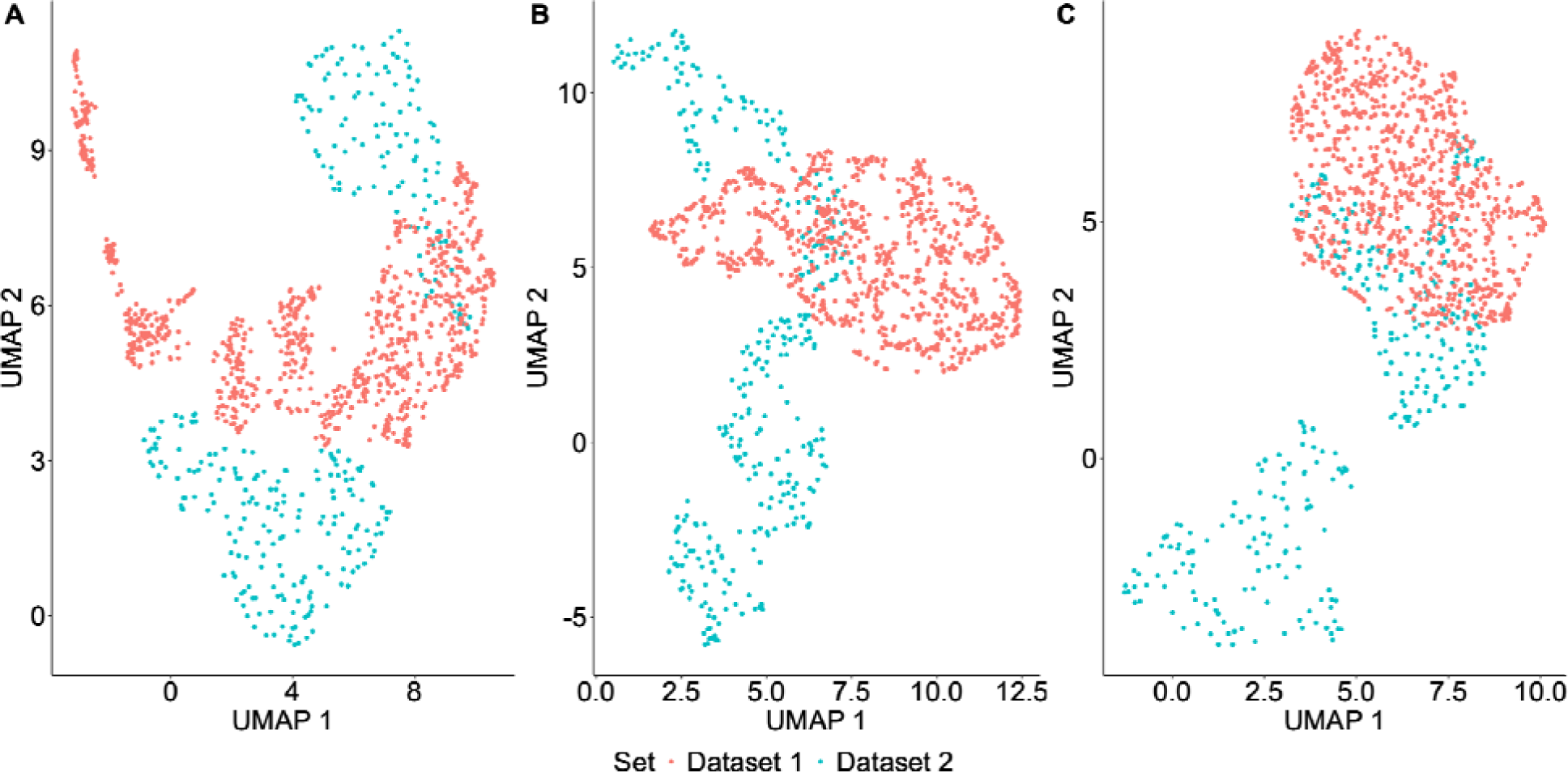
UMAPs obtained from input fragmentomics features illustrate batch effect between dataset 1 and dataset 2. (A) UMAP obtained from feature FLEN. (B) UMAP obtained from feature ND. (C) UMAP obtained from feature EM.

**Supplementary Figure 3.**
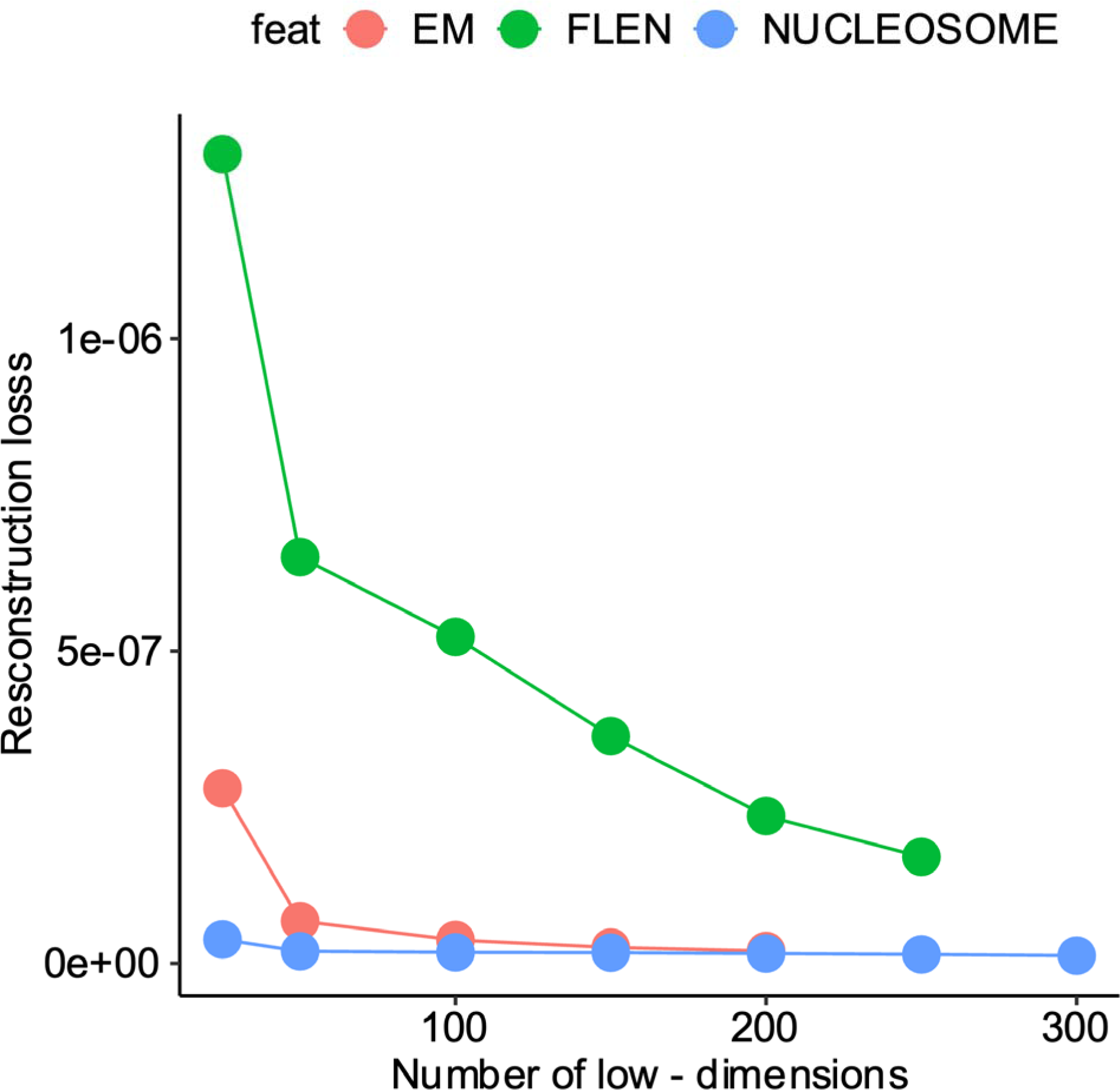
Reconstruction loss decreases when number of low-dimensions increases when transforming fragmentomics features with SNMF.

## List of abbreviations

cfDNA: Cell free DNA
NMF: Non-negative matrix factorization
SNMF: Supervised non-negative matrix factorization
FLEN: Fragment length distribution
ND: Distribution of distance of reads to nearest-nucleosome
EM: 4bp end-motif distribution
MM: Majorization – minimization algorithm
HCC: Hepatocellular carcinoma

